# Dually localized proteins found in both the apicoplast and mitochondrion utilize the Golgi-dependent pathway for apicoplast targeting in *Toxoplasma gondii*

**DOI:** 10.1101/2020.03.25.007476

**Authors:** Aparna Prasad, Pragati Mastud, Swati Patankar

## Abstract

Like other apicomplexan parasites, *Toxoplasma gondii* harbours a four-membraned endosymbiotic organelle - the apicoplast. Apicoplast proteins are nuclear-encoded and trafficked to the organelle through the endoplasmic reticulum (ER). From the ER to the apicoplast, two distinct protein trafficking pathways can be used. One such pathway is the cell’s secretory pathway involving the Golgi, while the other is a unique Golgi-independent pathway. Using different experimental approaches, many apicoplast proteins have been shown to utilize the Golgi-independent pathway, while a handful of reports show that a few proteins use the Golgi-dependent pathway. This has led to an emphasis towards the unique Golgi-independent pathway when apicoplast protein trafficking is discussed in the literature. Additionally, the molecular features that drive proteins to each pathway are not known. In this report, we systematically test eight apicoplast proteins, using a C-terminal HDEL sequence to assess the role of the Golgi in their transport. We demonstrate that dually localised proteins of the apicoplast and mitochondrion (*Tg*SOD2, *Tg*TPx1/2 and *Tg*ACN) are trafficked through the Golgi while proteins localised exclusively to the apicoplast are trafficked independent of the Golgi. Mutants of the dually localised proteins that localised exclusively to the apicoplast also showed trafficking through the Golgi. Phylogenetic analysis of *Tg*SOD2, *Tg*TPx1/2 and *Tg*ACN suggested that the evolutionary origins of *Tg*SOD2, *Tg*TPx1/2 lie in the mitochondrion while *Tg*ACN appears to have originated from the apicoplast. Collectively, with these results, for the first time, we establish that the driver of the Golgi-dependent trafficking route to the apicoplast is the dual localisation of the protein to the apicoplast and the mitochondrion.

## INTRODUCTION

Compartmentalisation is an indispensable feature in biological systems, especially for eukaryotic cells. Cellular compartments include endosymbiotic organelles, such as the mitochondria and chloroplasts, thought to have once been free-living prokaryotes that have since undergone an evolutionary transformation into the compartments that define the eukaryotic cell we know today. It is hypothesised that an important part of this transformation was gene transfer to the host nucleus (Herrmann, 1997) with the endosymbiont renouncing a large proportion of its genome and retaining only a limited set of genes. Hence, nuclear-encoded proteins, translated in the cytoplasm, contribute to the majority of the organellar proteome (Andersson et al., 2003; Richly, Chinnery & Leister, 2003; Richly & Leister, 2004). As endosymbiotic organelles exhibit a complex ‘orchestra’ of metabolic activity making them essential for cell survival, nuclear-encoded proteins that contribute to these specialised webs of interconnected metabolic pathways are targeted to the precise compartment, as if by design.

Certain endosymbiotic organelles are characteristic of distinct groups of organisms. For example, apicomplexa is a phylum of parasitic organisms including the human pathogens *Toxoplasma gondii* and *Plasmodium falciparum*; that harbour a secondary endosymbiotic plastid called the apicoplast (McFadden et al., 1996). The apicoplast is an organelle enclosed by four distinct membranes (Ferguson et al., 2005; Lemgruber et al., 2013), which, like the mitochondrion and chloroplast, has evolved specialised mechanisms for the import of nuclear-encoded proteins. Owing to a shared cyanobacterial ancestry, protein import into the inner two membranes of the apicoplast involves translocons of the Tic and Toc complexes, much like the chloroplast (van Dooren et al., 2008; Glaser et al., 2012; Sheiner et al., 2015). In addition to the apicoplast, apicomplexans also harbour another endosymbiotic organelle, the mitochondrion. This organelle, like other eukaryotic mitochondria, acquires newly synthesised proteins from the cytosol through translocons of the Tim and Tom complexes, proteins that appear to be derived from the outer membrane of the free-living prokaryote that was engulfed during endosymbiosis (van Dooren et al., 2016).

The presence of two outer membranes in the apicoplast differentiates the mechanism of protein import into the apicoplast from that of the chloroplast, where proteins with an N-terminal transit peptide are recognised by the Tic/Toc translocons (Perry & Keegstra, 1994; Ma et al., 1996; Kouranov & Schnell, 1997). Instead, for apicoplast proteins, a bipartite N-terminal signal sequence consists of a signal peptide preceding the transit peptide (Waller et al., 1998a). The signal peptide directs the protein to the endoplasmic reticulum (ER) and the transit peptide targets the protein to the apicoplast (Waller et al., 1998a). Uptake into the ER is the first step in the transport of proteins to the apicoplast and from the ER, the protein may take a Golgi-dependent or -independent pathway to reach the apicoplast. While a Golgi-dependent pathway can be paralleled to the usual secretory pathway (Chaudhari, Narayan & Patankar, 2012; Heiny et al., 2014), a Golgi-independent pathway is hypothesised to occur *via* transient contacts between the ER and apicoplast outermost membranes (Tomova et al., 2009) or through specialised vesicles between the ER and apicoplast (Heiny et al., 2014). Regardless of which of the two pathways is used by different proteins, the ER is central to the transport of proteins to the apicoplast and it is in the ER that the choice of route to the apicoplast is made.

There are many interesting hypotheses regarding the pathways taken by apicoplast proteins to reach their final destination. One such hypothesis revolves around the evolution of new modes of protein targeting to endosymbiotic organelles such as the apicoplast, in parallel to existing trafficking pathways (Reyes-Prieto, Weber & Bhattacharya, 2007; Li & Chiu, 2010). Here, it has been proposed that the Golgi-dependent secretory pathway is an ancient trafficking route (Reyes-Prieto, Weber & Bhattacharya, 2007), while the Golgi-independent pathway evolved later. Based on this evolutionary theory, it has been suggested that early in the evolution of endosymbiotic organelles, after transfer of genes to the nucleus, nuclear-encoded proteins used the existing secretory trafficking pathway to deliver proteins to the nascent endosymbionts (Bhaya & Grossman, 1991; McFadden, 1999). Some of these nuclear-encoded proteins would have contributed to the later evolution of new trafficking pathways, hence the Golgi-dependent pathway might have been used early during establishment of the endosymbiotic relationship and the Golgi-independent pathway might have evolved later. It is intriguing that for the apicoplast, there is evidence that the Golgi-dependent and - independent pathways of protein import exist to this day.

The Golgi-dependent pathway was first demonstrated in *P. falciparum* with glutathione peroxidase-like thioredoxin peroxidase (*Pf*TPx_Gl_), a protein localised to the apicoplast and the mitochondrion. Transport of *Pf*TPx_Gl_ to the apicoplast is disrupted in the presence of Brefeldin A (BFA), a fungal metabolite that inhibits retrograde transport between the Golgi and ER (Chaudhari, Narayan & Patankar, 2012). Additionally, a Golgi-independent pathway to the apicoplast is also known to exist in *P. falciparum*, where acyl carrier protein (*Pf*ACP), a protein that localises exclusively to the apicoplast, remains unaffected by the presence of BFA. This indicates that its transport is not dependent on the Golgi (Tonkin et al., 2006). However, in contrast to this finding, another report that used similar experimental strategies indicated that *Pf*ACP may use a Golgi-dependent route (Heiny et al., 2014). It is safe to say that in *P. falciparum*, the requirement of the Golgi for trafficking to the apicoplast is still an open area of research.

In *T. gondii, Tg*ACP, the homologue of *Pf*ACP that is also found exclusively in the apicoplast, has been shown to be unaffected by the presence of BFA (DeRocher et al., 2005). Further, addition of an ER retrieval sequence, HDEL (Hager et al., 1999), also did not affect the transport of *Tg*ACP to the apicoplast (DeRocher et al., 2005). As the HDEL motif results in ER localisation of proteins that transit through the Golgi, these data suggested that the Golgi has no role to play in *Tg*ACP transport (DeRocher et al., 2005). This indicates the existence of a Golgi-independent pathway in *T. gondii*. In contrast, a Golgi-dependent pathway to the apicoplast in *T. gondii* has yet to be proven with conclusive evidence although there are many clues indicating the likelihood of such a pathway. For example, for *Tg*SOD2, a dually localised protein of the apicoplast and mitochondrion in *T. gondii*, transport to the apicoplast is abolished in the presence of an HDEL sequence, while mitochondrial uptake is unaffected (Brydges, 2003). Unexpectedly, the presence of the HDEL sequence did not result in retrieval of *Tg*SOD2 to the ER. Clearly, more experiments are warranted for defining the Golgi-dependent pathway in *T. gondii*.

To clearly delineate features determining the choice of pathway employed by different proteins to reach the apicoplast, we analysed all apicoplast proteins with published experimental validation of their localisation in *T. gondii*. We tested the involvement of the Golgi in the transport of these proteins by assessing perturbation in the apicoplast localisation of these proteins in the presence of HDEL, the ER retrieval sequence in *T. gondii* (Hager et al., 1999). We provide conclusive evidence to establish that a Golgi-dependent pathway exists for proteins trafficked to the apicoplast in *T. gondii*. Our data also shows that proteins localised exclusively to the apicoplast employ a Golgi-independent pathway while dually localised proteins of the apicoplast and the mitochondrion traverse through the Golgi to reach the apicoplast. An analysis of the phylogeny and origins of these dually localised proteins indicates that these proteins could be mitochondrial in origin or have a cyanobacterial ancestry from the apicoplast. This report provides clarity on the ongoing controversy regarding Golgi-dependent and -independent trafficking to the apicoplast and also gives insights into the evolution of the N-terminal signals of proteins that target to this unique apicomplexan organelle.

## MATERIALS AND METHODS

### PLASMID CONSTRUCTION

#### *T. gondii* expression vectors for apicoplast proteins with C-terminal HA-tags

A list of extensively studied apicoplast-targeted proteins was generated (Table 1) and based on this data the following proteins were selected for experimental analysis: *Tg*TPx1/2 (TGME49_266120), *Tg*Der1_AP_(FJ976520.1), *Tg*ATrx2 (TGME49_310770), *Tg*ATrx1 (TGME49_312110), *Tg*PPP1 (JN053049.1), *Tg*Tic22 (TGME49_286050), *Tg*ACN (TGME49_226730) and *Tg*SOD2 (TGME49_316330). The genes encoding these proteins were cloned into pCTG-HA (Mastud & Patankar, 2019). This plasmid vector consists of pCTG-EGFP (van Dooren et al. 2008) which was modified to be devoid of GFP and contains a 1X HA tag. Total RNA was isolated from the parasites (wild type RH strain) using TRIsoln (Merck). This was used to generate cDNA using a gene specific reverse primer and RevertAid Reverse Transcriptase (Thermo Scientific). These genes were then amplified from cDNA using respective forward primers and reverse primers (Table 2). *Tg*TPx1/2, *Tg*Der1_AP_, *Tg*ATrx2, *Tg*ATrx1, *Tg*PPP1, *Tg*Tic22 and *Tg*SOD2 were cloned into pCTG-HA between MfeI and NdeI sites in pCTG-HA, generating C-terminally HA tagged genes of interest. Charged amino acid mutants, TPx1/2(R24A)-HA and SOD2(R12A)-HA, were generated by site-directed mutagenesis to replace the respective amino acids with an alanine using a gene specific primer against the plasmid containing the wild type constructs. The primers were designed with base pair mis-matches to replace an arginine with alanine. All clones were confirmed by sequencing.

**Table 1:** Experimentally validated data from literature on apicoplast proteins and dually targeted proteins of the apicoplast and mitochondrion in *Toxoplasma gondii*

| Localisation | Protein | SP-TP |  | Immuno-EM | Fractionation (membrane-association) | Function | References |
| --- | --- | --- | --- | --- | --- | --- | --- |
|  |  | Signal peptide Prediction | N-terminal cleavage |  |  |  |  |
| Apicoplast | ACP * | + | + | Lumen | Soluble | Metabolic pathway | (Waller et al. 1998; DeRocher et al. 2005; Glaser et al., 2012) |
|  | ATrx1 | + | + | Membrane associated | Soluble and Pellet | Redox/ apicoplast protein import | (DeRocher et al. 2008; Biddau et al. 2018 ) |
|  | ATrx2 | + | + | No data | No data | Redox | (Biddau et al. 2018) |
|  | Tic22 | + | + | No data | Soluble and Pellet | Apicoplast Protein import | (Glaser et al. 2012) |
|  | PPP1 | - | + | Peripheral | No data | Unknown function<sup>#</sup> | (Sheiner et al. 2011) |
|  | FtsH1 | - | + | Membrane | Pellet | <sup>^</sup>Metabolic pathway | (Karnataki et al. 2009) |
|  | Tic20 | + | + | Membrane | Pellet | Apicoplast protein import | (van Dooren et al. 2008a) |
|  | APT1 | - | - | Membrane | Pellet | Apicoplast ATP transport | (Karnataki et al. 2007)(Brooks et al., 2010) |
|  | Der1<sub>Ap</sub> | + | + | Membrane | No data | Apicoplast protein import | (Agrawal et al. 2009) |
| Apicoplast + Mitochondrion | TPx1/2 | + | + | No data | No data | Redox | (Pino et al., 2007; Mastud & Patankar, 2019) |
|  | SOD2 * | + | + | Lumen | No data | Redox | (Brydges et al., 2003; Pino et al., 2007) |
|  | ACN/IRP | + |  | No data | No data | Metabolic pathway | (Pino et al., 2007) |

Signal peptide prediction was done using SignalP 3.0 (Bendtsen et al., 2004) ‘+’ indicates positive prediction or experimentally validated N-terminal cleavage; ‘-’ indicates no prediction or no observed N-terminal cleavage. APT1 is the only exception with no signal prediction or N-terminal cleavage. ‘<sup>^</sup>’ indicates predicted function. <sup>#</sup>Periplastid protein 1 (PPP1) is shown to be important for parasite survival, apicoplast biogenesis and apicoplast protein import (Sheiner et al., 2011) \*ACP and SOD2 are the only two proteins that have been tested for the involvement of Golgi in the transport to the apicoplast, so far.

**Table 2:**
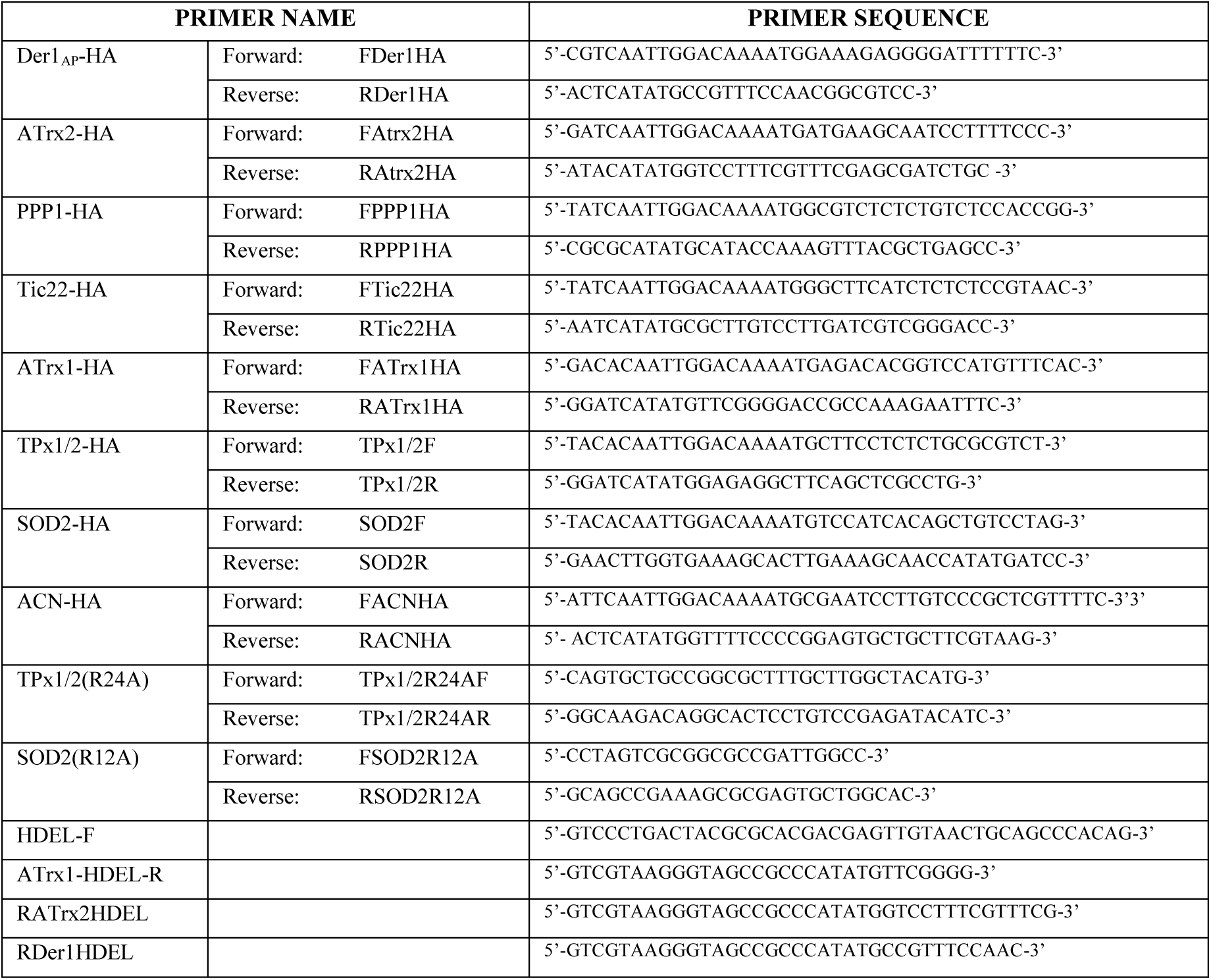
Primer sequences used for cloning genes

#### C-terminal HDEL constructs of apicoplast proteins

*Tg*TPx1/2-HA-HDEL and *Tg*SOD2-HA-HDEL were amplified as PCR products using TPx1/2F and SOD2R as forward primers and pCTGHDEL as a reverse primer using *Tg*TPX1/2-HA and *Tg*SOD2-HA as templates. The reverse primer, pCTGHDEL, contains the coding sequence for HDEL. The resultant PCR products were ligated between the MfeI and PstI sites of the vector pCTG-HA to generate the constructs. *Tg*ATrx1-HA-HDEL, *Tg*ATrx2-HA-HDEL and *Tg*Der1_AP_-HA-HDEL were generated by site directed mutagenesis using a common forward primer (HDEL-F) with a sequence coding for HDEL and sequence overlap with HA at the 5’ end of the primer and vector at the 3’ end. The reverse primer was specific to the gene at the 5’ end with a sequence overlap with the HA tag at the 3’ end (Atrx1-HDEL-R, RATrx2HDEL and RDer1HDEL). The constructs were confirmed by sequencing and *Tg*ATrx2-HA-HDEL was subsequently used as a vector for generating HDEL constructs after removing ATrx2 between MfeI and NdeI sites. *Tg*PPP1-HA-HDEL, *Tg*Tic22-HA-HDEL, were generated by cloning the respective gene in this vector between MfeI and NdeI sites. TPx1/2(R24A)-HA-HDEL and SOD2(R12A)-HA-HDEL were generated by site-directed mutagenesis to replace the respective amino acids with an alanine. All clones were confirmed by sequencing.

The plasmid pCTG-EGFP was obtained as a kind gift from Prof. Shobhona Sharma, Tata Institute of Fundamental Research, Mumbai.

#### *T. gondii* CULTURE, MAINTAINANCE AND TRANSFECTIONS

##### Culture and maintenance

*T. gondii* parasites (wild type RH strain) were cultured and maintained in primary human foreskin fibroblasts (HFF) (obtained from ATCC). HFF cells were maintained in T25 flasks in Dulbecco’s Modified Eagle Medium, DMEM (Gibco™, Mfr. No. 12100-046) containing 3.7 g/L sodium bicarbonate and 2.38 g/L HEPES, supplemented with 10% cosmic calf serum (Hyclone™) and 20 mg/L Gentamicin. Methods followed for maintenance and harvest of parasites were as described previously (Striepen & Soldati, 2007).

##### Transient transfections

Extracellular parasites (∼9-10 million) were re-suspended in 400 µl of DMEM devoid of serum and Gentamicin. DNA (50 µg) was mixed into this suspension in a 2 mM Bio-Rad GenePulser electroporation cuvette and pulsed at 1.5 kV, 50 Ω, 25 µF using a Bio-Rad GenePulser Xcell system. The transfected parasites were infected onto HFF cells and 24 hours post-transfection they were immuno-stained with appropriate antibodies.

*T. gondii* parasites expressing the protein PfTPx_Gl_47-EGFP (Narayan et al., 2018) were used as a parasite line expressing a GFP tagged ER marker into which *Tg*TPx1/2-HA-HDEL was transfected. The transfected parasites were infected onto HFF cells and 24 hours post-transfection they were immuno-stained with appropriate antibodies.

### IMMUNO-STAINING AND MICROSCOPY

Transiently transfected parasites were infected onto HFF cells growing in chamber slides (SPL Life Science Co.). 24 hours post-infection, the cells were fixed in 4% formaldehyde and 0.0075% glutaraldehyde in 1X PBS for 40 minutes. The cells were permeabilised with 0.25% TritonX-100 for 10 minutes. Blocking was done in 3% BSA in 1X PBS for one hour. This was followed by incubation with primary antibody in 3% BSA in 1X PBS for 2 hours at room temperature. The primary antibodies used were 1:1000 anti-HA Rabbit monoclonal antibody (C29F4; Cell Signaling Technology), 1:1000 anti-HA Mouse monoclonal antibody (Clone HA-7, mouse IgG1 Isotype; Sigma Aldrich) or 1:1000 anti-ACP antibodies (a kind gift from Prof. Dhanasekaran Shanmugam, NCL, Pune). The cells were washed thrice in 1X PBS and incubated with the appropriate secondary antibody in 3% BSA in 1X PBS for 1.5 hours at room temperature. The secondary antibodies used were 1:400 anti-rabbit Alexa 488 and 1:250 anti-mouse Alexa 594 from Invitrogen. The cells were washed thrice in 1X PBS, stained in DAPI (2µg/mL in 1X PBS) for 20 minutes at room temperature and washed twice with 1X PBS. The samples were mounted in ProLong Diamond Antifade mountant. Intracellular parasite mitochondrial staining was carried out using a previously described protocol (Salunke et al., 2018) with the fluorescent dye MitoTracker® red CMXRos (300 nM) from Invitrogen™.

The images were captured by a Zeiss LSM 780 NLO confocal microscope using a Plan-Apochromat 100X, 1.4 NA objective. Optical sections were captured at the interval of 0.44 µm. Maximum intensity projection images for these optical sections were obtained.

### IMAGE ANALYSIS

For determining colocalisation of the protein of interest with the organelle marker, at least 50 vacuoles of parasites were imaged at random for each construct. Thirty-five of these images for each construct were analysed; images that had intensities too high or too low to interpret results were not included in the analysis. For comparative analysis between wild type and C-terminal HDEL constructs, the imaging was done at the same settings. Image analysis was done using the software Imaris on unprocessed images. Using the ‘Coloc’ tool in Imaris, the volume of the image stacks occupied by the organelle marker was selected and tested for colocalisation of protein of interest with the marker. The resulting Pearson Correlation Coefficient (PCC) values were used to interpret colocalisation. To estimate the prevalence of a phenotype in the population the same images were used to identify the phenotype. Graphs depicting the population distribution were plotted using the GraphPad Prism version 8.0.1 (244) for Windows, GraphPad Software, La Jolla California USA. Representative images of each phenotype have been shown in the figures, which have been processed using ImageJ (Fiji). No non-linear adjustments such as changes to the gamma settings and curves were performed. All experiments were done in duplicates.

### ANALYSIS OF EVOLUTIONARY ORIGINS

To understand the evolutionary origins of *Tg*TPx1/2, *Tg*SOD2 and *Tg*ACN, homologues of these proteins were analysed from various groups of organisms. To differentiate between mitochondrial and apicoplast origins, plants and algae (apicoplast origins) and proteobacteria (mitochondrial origins) were included in this analysis. Apart from these, apicomplexans, other protists and eukaryotes were also included. *Tg*TPx1/2, *Tg*SOD2 and *Tg*ACN were subjected to domain prediction on PROSITE (de Castro et al., 2006). Proteins on the UniProt database (“UniProt,” 2019) that contain the respective domains and organisms that covered the desired groups of species were chosen. These proteins were confirmed for the presence of the respective domains by domain prediction on PROSITE and multiple sequence alignment using the MUSCLE algorithm (Edgar, 2004). Sequence similarity was tested using BLAST (Altschul et al., 1990).

Phylogenetic analyses were conducted using MEGA version X (Kumar et al., 2018). The multiple sequence alignment was done using the MUSCLE algorithm (integrated in MEGA) (Edgar, 2004). The phylogeny was inferred by using the Maximum Likelihood (ML) method and JTT matrix-based model (Jones, Taylor & Thornton, 1992). Partitions of branches that reproduced in less than 50% bootstrap replicates have not been shown. The percentage of replicate trees in which the associated taxa clustered together in the bootstrap test (500 replicates) (Felsenstein, 1985) are shown next to the branches. Initial tree(s) for the heuristic search were obtained automatically by applying Neighbor-Join and BioNJ algorithms to a matrix of pairwise distances estimated using a JTT model, and then selecting the topology with superior log likelihood value, as set to default settings in MEGA X (Kumar et al., 2018).

## RESULTS

### ANALYSIS OF FEATURES OF APICOPLAST-TARGETED PROTEINS

In order to address the question of molecular features that drive the choice of trafficking pathways of apicoplast proteins, the first step was to identify such features; these will be discussed individually. First, the localisation of a protein and its transport pathway to the destined organelle is largely driven by the N-terminus of the protein. Most proteins destined to the apicoplast have bipartite signalling sequences at their N-termini (Waller et al., 1998b), directing them to the apicoplast, while proteins destined to the mitochondrion have mitochondrial targeting sequences mediating their uptake into the mitochondrion (Hay, Böhni & Gasser, 1984; Douglas, McCammon & Vassarotti, 1986). A protein targeted to both the mitochondrion and the apicoplast may have an ambiguous signal sequence that has components of both the organellar signal sequences (Chew et al., 2003; Mastud & Patankar, 2019). These N-termini have evolved to dictate the localisation and consequently the pathway by which the protein reaches the apicoplast. Therefore, we recorded whether a protein is localised to the apicoplast alone or dually targeted to both the apicoplast and mitochondrion.

Apart from the N-terminal sequence, other features of the protein may influence its localisation and choice of transport pathway. These factors include: N-terminal cleavage, localisation within the apicoplast, association with the membrane and function of the protein. For example, if a protein is cleaved at the N-terminus, this might expose the binding site for translocons in the ER that route the processed protein via the Golgi-independent pathway. In contrast, a lack of cleavage of the N-terminus might hide such binding sites, while still allowing binding to Golgi-dependent trafficking proteins such as SNAREs (SNAP Receptor proteins).

The location of an apicoplast protein in one or more of the four membranes, or in the intermembrane spaces, might also affect the choice of trafficking route. Here, it could be hypothesized that proteins found in the outmost membrane might reach that membrane via a different pathway compared to proteins that are found in the lumen. Finally, protein function might require post-translational modifications, such as glycosylation, that take place in the Golgi, leading to some proteins being trafficked via the secretory pathway.

Experimentally validated data of these factors for apicoplast proteins and dually targeted proteins of the apicoplast and mitochondrion have been collated in Table 1. This table includes only those proteins for which the aforementioned data are available in the literature. Based on their localisation, the proteins in Table 1 have been broadly categorised as apicoplast proteins and dually localised proteins (apicoplast + mitochondrion). The presence of an N-terminal signal peptide and its cleavage are important features of apicoplast proteins; the signal peptide directs the protein to the ER and cleavage exposes the transit peptide important for uptake into the apicoplast. Table 1 indicates that these features are present in dually localised proteins as well and seem to be essential features of proteins directed to the apicoplast irrespective of their localisation.

Proteins shortlisted in Table 1 can also be categorised based on their membrane association. Apicoplast proteins are either luminal proteins, membrane proteins or have been observed by immuno-EM to be present in the intermembrane space, but closely associated with the membrane, for example *Tg*ATrx1 and *Tg*PPP1 (DeRocher et al., 2008; Sheiner et al., 2011; Biddau et al., 2018). Among the dually localised proteins, only *Tg*SOD2 is reported to be a luminal protein while there are indications that *Tg*TPx1/2 may be associated with the membrane (unpublished data). *Tg*ACN/IRP has not been tested.

Proteins in Table 1 have also been broadly categorised based on their function as redox proteins, proteins involved in metabolic pathways or proteins involved in protein transport. This categorisation is based on both experimental validations as well as hypothesised functions. The dataset of apicoplast proteins contains those from all three categories as expected. Two of the dually localised proteins, *Tg*SOD2 and *Tg*TPx1/2, are redox proteins while *Tg*ACN/IRP is hypothesised to be a component of a metabolic pathway (Pino et al., 2007). Although the dataset of dually localised proteins is very limited it is interesting to note that two of the three proteins shared between the apicoplast and the mitochondrion are redox proteins.

As described before, transport of proteins to the apicoplast in *T. gondii* can be both Golgi-dependent and - independent. Differences and commonalities of these features among all the proteins in Table 1 might provide clues to factors that influence the choice of route to the apicoplast. However, the involvement of Golgi in transport to the apicoplast has been tested for only two proteins, so far (indicated by * in Table 1). The transport of an exclusively apicoplast resident acyl carrier protein (*Tg*ACP) to the apicoplast is unaltered in the presence of a C-terminal ER retention sequence—HDEL (DeRocher et al., 2005). *Tg*SOD2, a dually targeted protein of the mitochondrion and apicoplast, does not localise to the apicoplast in the presence of this motif (Brydges, 2003), however does not show ER retention either. Evidently, a comprehensive analysis of apicoplast proteins is required to infer the route that they employ to the apicoplast. Hence, the exclusively apicoplast proteins (*Tg*ATrx1, *Tg*ATrx2, *Tg*Tic22, *Tg*PPP1, and *Tg*Der1_AP_) and the dually localised proteins of the mitochondrion and apicoplast (*Tg*TPx1/2, *Tg*SOD2 and *Tg*ACN) were chosen to test involvement of Golgi in their transport to the apicoplast and further understand factors that might influence this choice of route.

### LOCALISATION OF APICOPLAST PROTEINS IS UNAFFECTED BY THE PRESENCE OF A C-TERMINAL HDEL SEQUENCE

To test the involvement of the Golgi in the transport of proteins to the apicoplast we added an HDEL sequence to the C-termini of the proteins of interest. An HDEL sequence in *T. gondii* is an ER-retention signal (Hager et al., 1999), much like the SDEL motif in *Plasmodium falciparum* (Kumar & Zheng, 1992) or the KDEL motif in mammalian cells (Munro & Pelham, 1987). These sequences function through interactions with HDEL receptors in the Golgi, resulting in the recycling of HDEL-bearing proteins back to the ER (Lewis, Sweet & Pelham, 1990; Semenza et al., 1990). Therefore, it would be expected that if a protein traverses the Golgi, the addition of an HDEL sequence would hamper the protein’s localisation to the target organelle and instead result in accumulation in the ER. This strategy has been used in *T. gondii* to test the involvement of the Golgi in the transport of proteins to the apicoplast (DeRocher et al., 2005). It is important to note that this strategy is limited to proteins whose C-termini face the lumen of the ER/Golgi, since the HDEL receptors themselves face the lumen (Scheel & Pelham, 1998). Hence, proteins whose C-termini are predicted or proven to be facing the cytosol were not included in this study. After this filtration step, C-terminal HA tag constructs of apicoplast proteins (*Tg*ATrx1, *Tg*ATrx2, *Tg*Tic22, *Tg*PPP1, *Tg*Tic20 and *Tg*Der1_AP_) were generated. An HDEL sequence was then added to these constructs. Expression plasmids for each gene, with and without the HDEL tag were transiently transfected into *T. gondii* parasites and the localisation of the proteins assessed by immunofluorescence using anti-HA antibodies.

Previously published data has indicated that the proteins *Tg*Der1_AP_, *Tg*ATrx1, *Tg*ATrx2, *Tg*PPP1 and *Tg*Tic22 are apicoplast resident proteins (DeRocher et al., 2008; Agrawal et al., 2009; Sheiner et al., 2011; Glaser et al., 2012; Biddau et al., 2018). Consistent with this data, C-terminal HA fusions of these proteins were localised to the apicoplast, as seen by colocalisation with the apicoplast marker ACP (Fig.1). Interestingly, the addition of an HDEL sequence did not change the apicoplast localisation of these proteins, indicating that these proteins do not traverse the Golgi during transit to the apicoplast (Fig.1). The lack of disruption in localisation with an HDEL sequence was independent of the spatial distribution within the apicoplast, i.e. whether the protein is found in the lumen, different membranes or inter-membrane compartments. Additionally, the apicoplast localisation of both membrane-associated and soluble proteins was unaffected by the addition of the HDEL sequence. Hence, the parameters shown in Table 1 do not influence the choice of route to the apicoplast, in this case a Golgi-independent route. The one common factor among all these proteins is that they are localised to the apicoplast alone. This raises the question of whether dual localisation is the key factor that defines whether proteins employ a Golgi-dependent pathway for trafficking to the apicoplast.

**Fig. 1:**
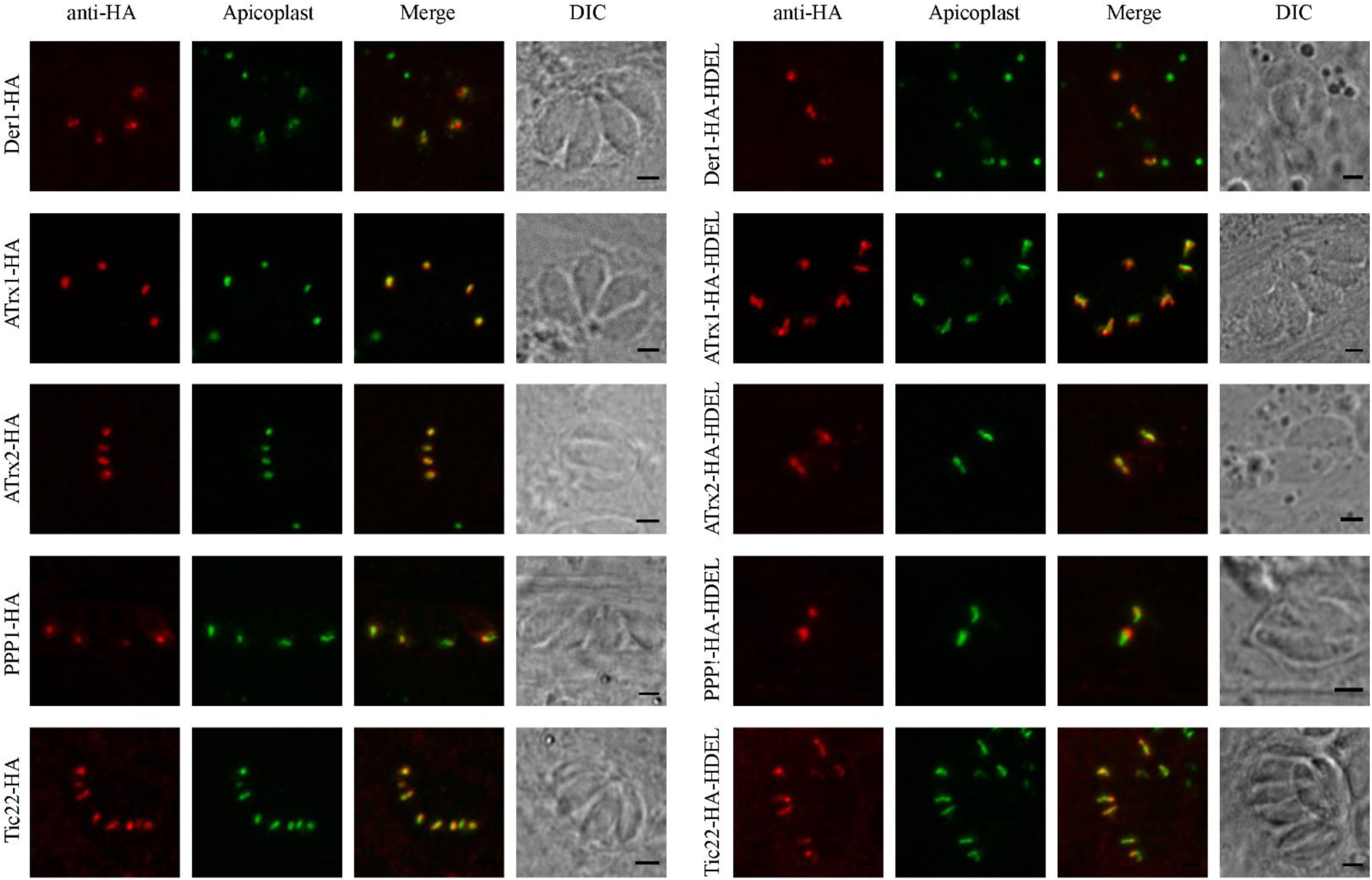
Localisation of apicoplast proteins with and without a C-terminal HDEL sequence Each protein is tagged with a C-terminal HA tag, shown in red. The apicoplast is stained with anti-ACP antibodies, as an apicoplast marker, in green. The panels to the left are the HA tagged apicoplast proteins, on the right are the corresponding constructs with a C-terminal HDEL.

### TRANSPORT OF DUALLY LOCALISED PROTEINS TO THE APICOPLAST IS PERTURBED BY A C-TERMINAL HDEL MOTIF

*Tg*TPx1/2, *Tg*SOD2 and *Tg*ACN/IRP are dually localised proteins in *T. gondii* targeted to both the apicoplast and the mitochondrion (Brydges, 2003; Pino et al., 2007). To test whether these proteins traverse the Golgi, C-terminal HA-tagged constructs of their cDNAs were generated and an HDEL sequence was then added to the C-terminus of each protein. Upon transfection of the plasmids without the HDEL sequence, *Tg*SOD2-HA was seen to localise both to the apicoplast and mitochondrion (Fig. 2B), as has been reported previously by Pino et al. A C-terminal HDEL sequence led to a complete loss of localisation to the apicoplast, as seen in Fig.2B, whereas targeting to the mitochondrion remained unaffected. No retention of the protein was observed in the ER. This is consistent with previously reported data that suggests that an HDEL motif abolishes only apicoplast targeting of *Tg*SOD2 (Brydges, 2003).

**Fig. 2:**
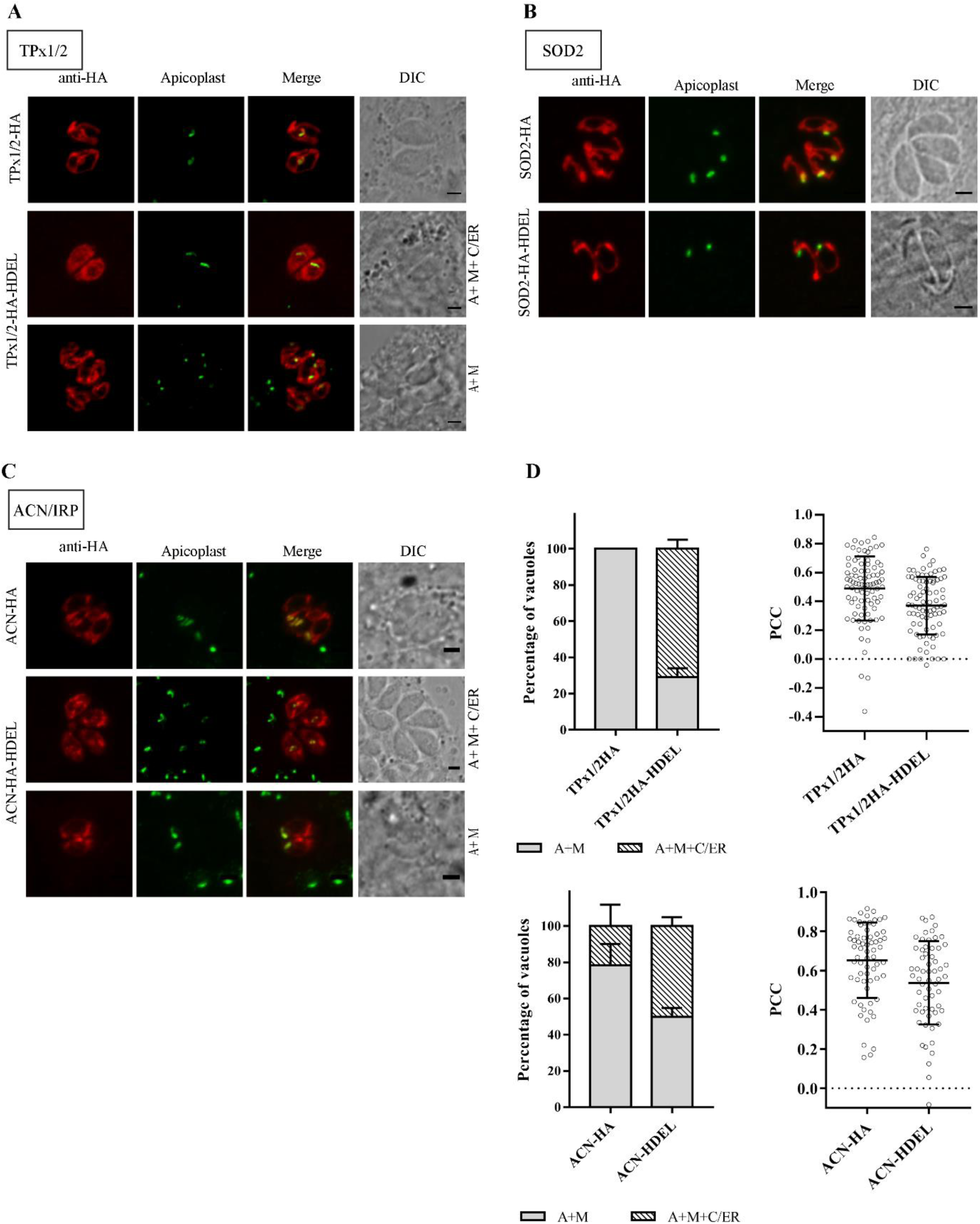
Effect of an HDEL sequence on the localisation of dually localised proteins to the apicoplast. Each protein is tagged with a C-terminal HA tag, shown in red. The apicoplast is stained with anti-ACP antibodies, as an apicoplast marker, in green. The cells have been observed to exhibit two phenotypes; A+M: Apicoplast and mitochondria, A+M+C/ER: Apicoplast and mitochondrion with a cytosol and ER like appearance. Each of the panels represent phenotypes of A) *Tg*TPx1/2-HA and *Tg*TPx1/2-HA-HDEL, B) *Tg*SOD2-HA and *Tg*SOD2-HA-HDEL, C) *Tg*ACN/IRP-HA and *Tg*ACN/IRP-HA-HDEL. D) Represents two graphs: the percentage of population exhibiting each phenotype of A+M and A+M+C/ER, and the mean PCC values for colocalisation with the apicoplast marker, of *Tg*TPx1/2 and *Tg*ACN/IRP with and without an HDEL sequence.

*Tg*TPx1/2-HA also localised to the apicoplast and the mitochondrion, consistent with previously published data (Pino et al., 2007; Mastud & Patankar, 2019). Interestingly, with the presence of a C-terminal HDEL motif, for ∼70% of the population, the protein localised to the apicoplast and was also perinuclear and spread across the cell (Fig.2A). This perinuclear staining colocalised with an ER marker (S1) and also appears to be diffused across the cytoplasm the cell. Targeting of this protein to the mitochondrion was undisturbed. As multiple phenotypes were observed in this experiment, a quantitative analysis of colocalisation was performed. As compared to *Tg*TPx1/2-HA, a significant decrease was observed in the average PCC value for colocalisation of TgTPx1/2-HA-HDEL with the apicoplast marker (Fig.2D). This indicates that the transport of *Tg*TPx1/2 to the apicoplast is perturbed in the presence of an HDEL sequence. It is noteworthy that, unlike *Tg*TPx1/2-HA-HDEL, *Tg*SOD2-HA-HDEL did not show any accumulation of protein in the ER despite containing an ER-retention signal.

*Tg*ACN/IRP has been previously reported to be localised to the apicoplast and the mitochondrion (Pino et al., 2007). Consistent with this report, we also found that *Tg*ACN-HA localised to both the apicoplast and mitochondrion (Fig.2C). Similar to the results observed for *Tg*TPx1/2, upon addition of the HDEL sequence, *Tg*ACN-HA-HDEL showed a perinuclear spread along with localisation to the apicoplast and the mitochondrion (Fig.2C), similar to that of *Tg*TPx1/2-HA-HDEL. This phenotype was seen in ∼50% of the population as compared to ∼21% in *Tg*ACN-HA (Fig.2D). *Tg*ACN-HA-HDEL showed a significant decrease in the PCC values for colocalisation with an apicoplast marker, as compared to *Tg*ACN-HA (Fig.2D) thus, indicating a decrease in apicoplast localisation and an increase in ER localisation upon the addition of an HDEL sequence. Taken together, these results suggest that proteins dually localised to the apicoplast and mitochondrion are trafficked via a Golgi-dependent route.

### A C-TERMINAL HDEL SEQUENCE INCREASES RETENTION OF *Tg*TPx1/2(R24A) AND *Tg*SOD2(R12A) IN THE ER

Data shown so far points to the dual localisation of proteins to the apicoplast and the mitochondrion being the factor that directs these proteins through the Golgi. In *Tg*TPx1/2 and *Tg*SOD2 this dual localisation is known to be driven by ambiguous N-terminal signal sequences. Hence, we next tested the effect of abolishing the dual localisation of these proteins on their choice of route to the apicoplast. In the case of *Tg*TPx1/2, the presence of an immunofluorescence signal both at the mitochondrion and the apicoplast made it difficult to definitively establish the identity of the perinuclear spread. Contrastingly, for *Tg*SOD2, an HDEL sequence abolished the transport to the apicoplast but did not result in ER-retention as expected. Therefore, upon the addition of an HDEL sequence, *Tg*TPx1/2 and *Tg*SOD2 presented two very different phenotypes of disruption in transport to the apicoplast, while their mitochondrial localisation was unperturbed.

In order to test whether dual localisation was the key factor in the choice of trafficking pathway and to eliminate the prominent signal from the mitochondrion, the proteins were mutated to localise exclusively to the apicoplast and were tested for involvement of the Golgi in the transport to the apicoplast. This was done by mutating the single positively charged amino acid in the N-terminus of the protein. Previous experiments on dually localised proteins *Tg*TPx1/2 and *Tg*SOD2 have demonstrated that mutating the single positively charged amino acid of the ambiguous N-terminal sequence completely abolished mitochondrial targeting (Brydges, 2003; Mastud & Patankar, 2019).

The single positively charged amino acid in the N-terminus of *Tg*TPx1/2, an arginine at position 24, was replaced with alanine to generate the construct *Tg*TPx1/2(R24A)-HA. Consistent with published data (Mastud & Patankar, 2019), this construct colocalised with the apicoplast marker ACP and showed no mitochondrial localisation (Fig.3A). In some cases, in addition to apicoplast localisation, the protein also showed localisation to the ER (Mastud & Patankar, 2019). An arginine at position 12 in the N-terminus of *Tg*SOD2 was replaced with an alanine to generate *Tg*SOD2(R12A)-HA. Consistent with previously published data (Brydges, 2003), *Tg*SOD2(R12A)-HA localised to the apicoplast. As seen in Fig.3D, this protein colocalised with the apicoplast marker ACP and no mitochondrial localisation was seen. In addition to apicoplast localisation, this protein showed localisation similar to *Tg*TPx1/2(R24A)-HA in the ER, in some rosettes.

In this study, about 60% of the population in the case of *Tg*TPx1/2(R24A)-HA and 21% in the case of *Tg*SOD2(R12A)-HA showed mixed phenotypes of localisation in both the apicoplast and the ER (Fig.3C, F). The remaining population of rosettes showed exclusive apicoplast localisation for both the proteins. Brydges and Carruthers have previously reported that on addition of an HDEL sequence, *Tg*SOD2(R12A) is localised to the apicoplast and also retained in the ER. Similar to this study, Brydges and Carruthers used a transiently transfected overexpression system, while they used GFP as a reporter gene unlike this study, where all the constructs are HA-tagged.

To investigate the involvement of Golgi in the transport of *Tg*TPx1/2 and *Tg*SOD2 to the apicoplast, an HDEL motif was added to C-terminus of *Tg*TPx1/2(R24A)-HA and *Tg*SOD2(R12A)-HA and transiently transfected parasites were analysed for localisation of the proteins. It was observed that both these constructs, *Tg*SOD2(R12A)-HA with and without a C-terminal HDEL, showed two phenotypes: localisation to the apicoplast alone and apicoplast localisation along with ER (Fig.3D, E). On comparing the distribution of these phenotypes within the population, it was observed that the addition of an HDEL sequence significantly increased the ER localisation of *Tg*SOD(R12A) (Fig.3F). While 21% of the population localised to the ER along with the apicoplast, in the case of *Tg*SOD2(R12A)-HA, the proportion of this phenotype increased to 69% upon addition of an HDEL sequence (Fig. 3F)

**Fig. 3:**
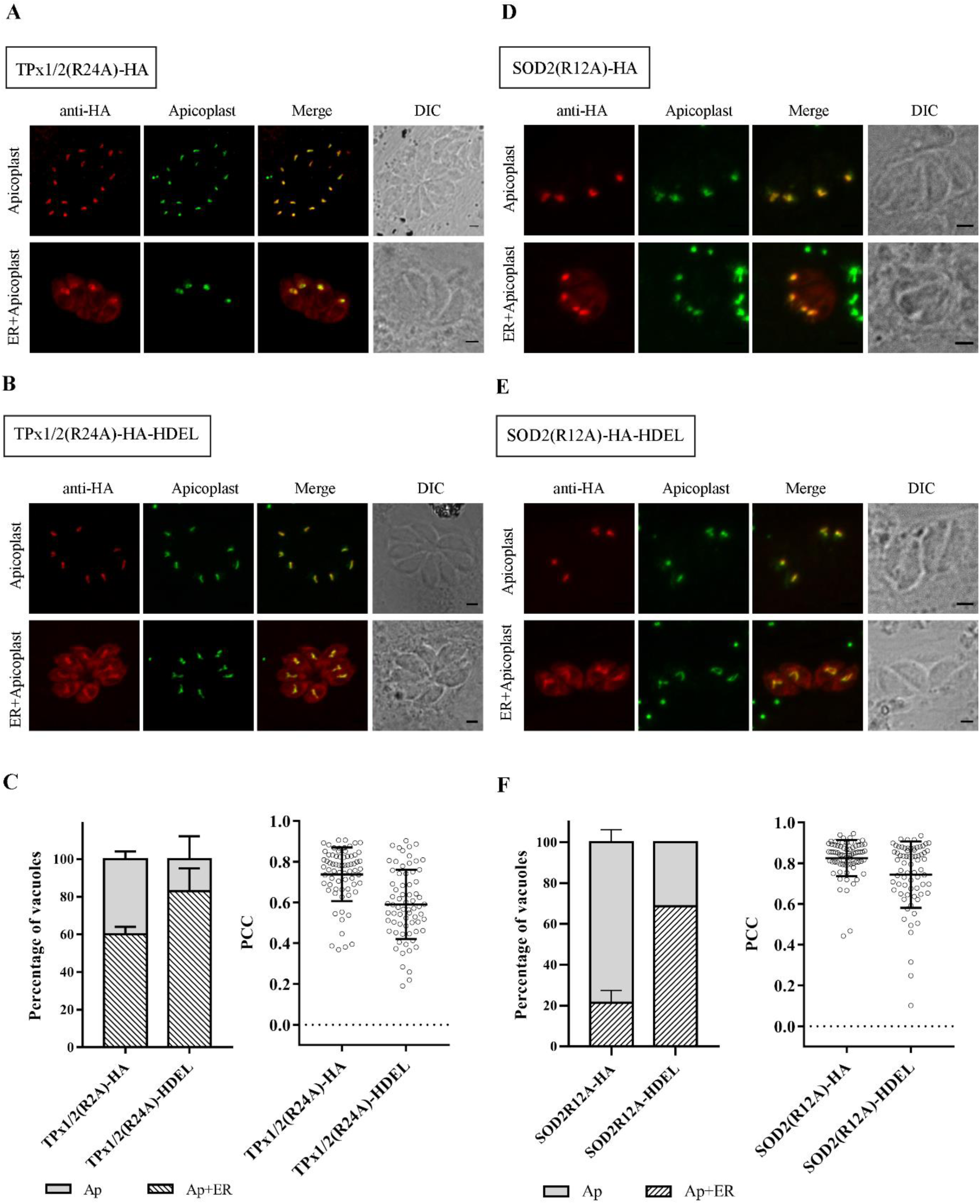
Effect of single charged residue mutations of the N-terminus on the localisation of dually localised proteins to the apicoplast and the effect of an HDEL sequence on these constructs. Each protein is tagged with a C-terminal HA tag, shown in red. The apicoplast is stained with anti-ACP antibodies, as an apicoplast marker, in green. The cells have been observed to exhibit two phenotypes; Ap: Apicoplast, Ap+ER: Apicoplast and ER. Each of the following panel represents the phenotype of A) *Tg*TPx1/2(R24A)-HA, B) *Tg*TPx1/2(R24A)-HA-HDEL, D) *Tg*SOD2(R12A)-HA, E) *Tg*SOD2(R12A)-HA-HDEL. C) and F) represent two graphs each: the percentage of population exhibiting each phenotype of Ap and Ap+ER, and the mean PCC values for colocalisation with apicoplast marker, of *Tg*TPx1/2(R24A) and *Tg*SOD2(R12A) with and without an HDEL sequence.

Similarly, two phenotypes of localisation, to the apicoplast alone and localisation to both the apicoplast and ER, were observed in a population of parasites transiently transfected with *Tg*TPx1/2(R24A)-HA and *Tg*TPx1/2(R24A)-HA-HDEL (Fig.3A, B). Again, an HDEL sequence markedly increased the population demonstrating an ER localisation along with apicoplast. The proportion of the population where the protein was localised to the apicoplast and the ER, with *Tg*TPx1/2(R24A)-HA, was 60%. With a C-terminal HDEL sequence, this proportion increased to 83% (Fig.3C).

Collectively, these data show that a C-terminal HDEL motif results in ER-retention of the *Tg*TPx1/2-HA protein and a loss of apicoplast localisation of the *Tg*SOD2 protein, while the HDEL motif increases the retention of both *Tg*TPx1/2(R24A) and *Tg*SOD(R12A) in the ER. This confirms that for these two proteins, whether dually localised as wild type proteins, or localised to the apicoplast alone, as mutant proteins, trafficking to the apicoplast is through a Golgi-dependent pathway.

### DUALLY LOCALISED PROTEINS, TRAFFICKED VIA THE GOLGI, HAVE MULTIPLE EVOLUTIONARY ORIGINS

As mentioned previously, it has been proposed that the ER-Golgi pathway is an ancient trafficking route, already in existence when the nascent endosymbiont was acquired by the emerging plastid-harbouring organism. During this period, transfer of genes from the genome of the endosymbiont to the host nuclear genome was presumably taking place. Finally, these genes acquired signals that would direct their transport back to the apicoplast, therefore, nuclear-encoded apicoplast proteins might be expected to show an evolutionary ancestry closer to plants. Rather than acquiring transit peptides from plant origins, these sequences have been proposed to be acquired from apicomplexan exons due to their rather loose sequence requirements (Tonkin et al., 2008), while the signal peptides might have been acquired later and fused with the existing transit peptides.

How then, might proteins that are dually localised to both the apicoplast and mitochondrion have evolved? Intriguingly, both mitochondrial targeting signals and transit peptides have similar features, including positively charged residues (DeRocher et al., 2000). Therefore, one might speculate that existing mitochondrial proteins acquired a signal peptide and were diverted to the ER and subsequently the apicoplast. Alternatively, apicoplast proteins whose genes had been transferred to the host nucleus might have possessed transit peptides that closely resembled mitochondrial targeting signals and then later acquired signal peptides. Evidence in favour of the second possibility lies in the case of the S9 ribosomal protein which is targeted to the apicoplast (Waller et al., 1998a; DeRocher et al., 2000). A phylogenetic analysis of S9 from apicomplexans reveals that it is more closely related to plant S9 proteins, suggesting that the origins of the gene encoding the S9 protein appear to lie with the endosymbiotic ancestor (Waller et al., 1998a). Deletion analysis of the N-terminal targeting sequences of the *T. gondii* S9 protein reveals that upon removal of the signal peptide, the transit peptide directs this apicoplast protein to the mitochondrion (DeRocher et al., 2000), indicating that the apicoplast transit peptide carries information that can be recognised by the mitochondrial Tim/Tom translocons after synthesis in the cytosol.

What then, are the connections between N-terminal targeting signals and trafficking routes? Data shown in this report reveal that proteins that are dually targeted to both the apicoplast and mitochondrion employ the ER-Golgi pathway for trafficking to the apicoplast, and this pathway has been proposed to be an ancient pathway, present before the Golgi-independent pathway. Perhaps the three dually localised proteins (*Tg*SOD2, *Tg*TPx1/2 and *Tg*ACN/IRP) were mitochondrial proteins encoded by the host genome that were later co-opted into the apicoplast by acquisition of signal peptides. They used the available trafficking pathway at the time, the Golgi-dependent route, which continues to this day. We tested this hypothesis by carrying out phylogenetic analyses of the three proteins, with the expectation that, despite being apicoplast-targeted proteins, they would lie closer to bacteria and mitochondrial homologues, rather than plants on the phylogenetic trees. Indeed, for one of the three proteins, *Tg*SOD2, this is already known to be the case (Sienkiewicz et al., 2004).

Homologues of *Tg*TPx1/2 chosen for this analysis have a peroxidase functional domain and interestingly, a large number of these eukaryotic homologues with high sequence homology to *Tg*TPx1/2 were mitochondrial proteins (based on information available on UniProt). Consistent with this preliminary observation, phylogenetic analysis of *Tg*TPx1/2 with its homologues selected in related groups of organisms clearly indicates a divergence from plants and algae (Fig.4). *Tg*TPx1/2 is seen to group with apicomplexans, as expected and further with other eukaryotes like fungi. The protein is more closely related to bacteria than to plants indicating that it could be a protein of mitochondrial origin that gained function at the apicoplast. This is a similar result to that seen for *Tg*SOD2 which has been reported to be related closely to protozoans and bacteria, compared to chloroplasts or chlorophytes, indicating its mitochondrial origins (Sienkiewicz et al., 2004). Interestingly, both these proteins have anti-oxidant functions. In contrast, when a similar analysis was performed for *Tg*ACN, it was observed that this protein is more closely associated to plant ACN proteins (both chloroplast and mitochondrial ACNs) than it is to other eukaryotic or bacterial ACNs (Fig.4). Thus, *Tg*ACN’s evolutionary origins are similar to other plant-derived apicoplast proteins like S9.

**Fig. 4:**
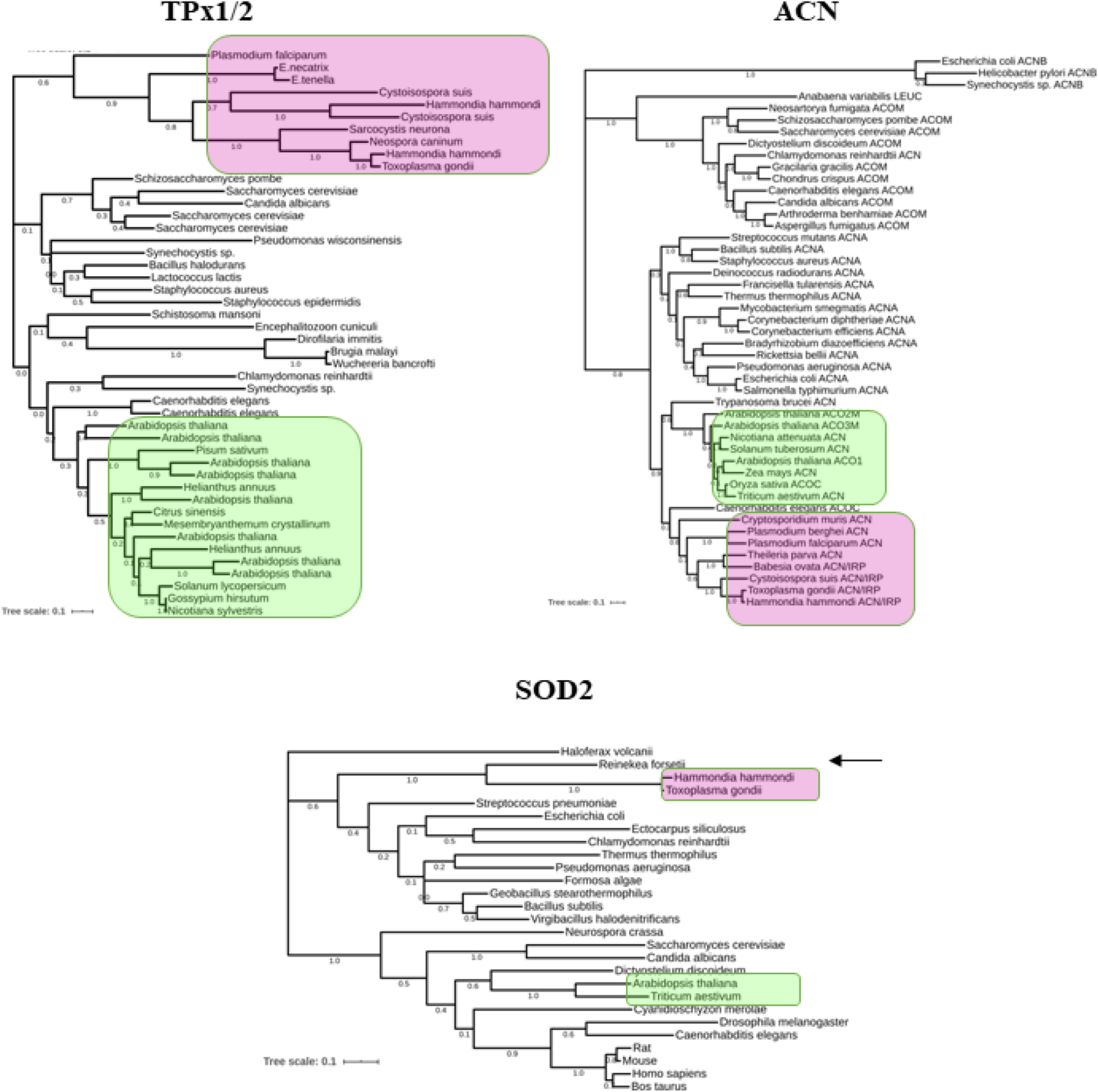
The evolutionary origins of *Tg*TPx1/2, *Tg*ACN and *Tg*SOD2. The phylogenetic relations of these proteins to their homologues in Bacteria, plants, algae and other eukaryotes has been established. The trees were generated by using the Maximum Likelihood method and JTT matrix-based model. In a bootstrap analysis of 500 repeats, the percentage of trees in which the respective taxa clustered together is shown. The tree is drawn to scale, with branch lengths measured in the number of substitutions per site. *Tg*TPx1/2 is seen to be closely related to bacteria and fungi as opposed to plants. *Tg*SOD2 is clearly seen to share ancestry with a proteobacterium *Reinekea forseti* (highlighted with an arrow). While, *Tg*ACN seems to be more closely related to plants than it is to proteobacteria or other eukaryotes. Plant species are highlighted in green while those of phylum Apicomplexa are highlighted in purple.

Phylogenetic analysis of the three dually targeted proteins indicates that *Tg*TPx1/2 and *Tg*SOD2, both redox proteins, may have originated from the mitochondrion and gained a function at the apicoplast, while *Tg*ACN may have originated from the apicoplast and is now functional at the mitochondrion as well. Hence, proteins that were encoded by the ancestral host genome as well as those acquired from the nascent endosymbiont appear to have hitchhiked on the host’s secretory pathway, using the ER-Golgi route for trafficking to the apicoplast.

## DISCUSSION

Proteins can reach the apicoplast *via* two separate pathways originating at the ER. This has been shown in *T. gondii* and a related apicomplexan parasite, *P. falciparum* (Waller et al., 2000; DeRocher et al., 2005; Tonkin et al., 2006; Chaudhari, Narayan & Patankar, 2012; Heiny et al., 2014; Chaudhari et al., 2017), with some proteins using a Golgi-dependent pathway and others using a Golgi-independent pathway. The determinants on each protein that determine the choice between these pathways are unclear and even more confusingly, in the case of *P. falciparum* ACP, two studies report opposite results, with one showing a Golgi-dependent pathway (Heiny et al., 2014) and the other showing a Golgi-independent pathway (Tonkin et al., 2006). In this study we clearly show the existence of these two pathways in *T. gondii* and determine the features that affect the choice of pathway. For the first time, this study establishes that the dual localisation of proteins, to both the apicoplast and mitochondrion, drives proteins through the Golgi while proteins localising solely to the apicoplast favour a Golgi-independent pathway.

### APICOPLAST PROTEINS ARE TRAFFICKED TO THE APICOPLAST INDEPENDENT OF THE GOLGI

Using the well-established tool of the addition of an ER-retention sequence to determine whether a particular protein traverses the Golgi, we show that proteins exclusively localised to the apicoplast are transported independently of the Golgi. This choice of pathway was seen to be independent of the protein localisation within the apicoplast, membrane association, signal processing and function— all factors that were hypothesised to potentially affect this choice.

Six apicoplast proteins were systematically tested in this study: *Tg*ATrx1, *Tg*ATrx2, *Tg*Tic22, *Tg*PPP1, *Tg*Tic20 and *Tg*Der1_AP_. Their transport to the apicoplast remains unhindered in the presence of an HDEL sequence. This is consistent with previously reported data on the luminal protein *Tg*ACP, whose transport to the apicoplast was not disturbed in the presence of an HDEL sequence (DeRocher et al., 2005). However, the remaining five proteins had not been directly tested before this study; instead, indirect evidence suggested a lack of Golgi involvement. This evidence included the observation that membrane proteins of the apicoplast like *Tg*ATrx1, *Tg*FtsH1 and *Tg*APT1 persist in large vesicles around the apicoplast (Karnataki et al., 2007b,a; DeRocher et al., 2008) hypothesised to be involved in their transport, after plastid loss and disruption of the Golgi. This indicated that the transport of these proteins may be independent of the Golgi. We have tested *Tg*ATrx1 and demonstrated that its transport is indeed independent of the Golgi, concurrent with previously reported observations. Therefore, the large vesicles reported previously appear to be involved in the trafficking of *Tg*Atrx1 directly from the ER to the apicoplast. *Tg*APT1 and *Tg*FtsH1, which are membrane proteins with cytosolic C-termini, could not be tested due to the technical constraints of using an HDEL sequence. Further evidence for the ER-apicoplast vesicles being distinct from secretory vesicles lies in a study in *P. falciparum* that showed, using chemical inhibition of G-protein coupled vesicular transport, that these vesicles were not affected by treatment and are different from the vesicles going to the Golgi (Chaudhari et al., 2017). Isolation and characterisation of the protein and lipid repertoire of these large vesicles would shed light on mechanisms of ER-apicoplast transport in *T. gondii*.

For a protein to be targeted to the apicoplast directly from the ER, and independent of the Golgi, the transit peptide may be a key player. Receptors specific to the transit peptide, in the ER, might be involved in the recognition of apicoplast proteins and the consequent transport to the apicoplast. Higher affinity of these transit peptides to their receptors as compared to the transit peptides of dually localised proteins could be a factor mechanistically driving apicoplast proteins along the Golgi-independent pathway.

### DUALLY LOCALISED PROTEINS OF THE APICOPLAST AND MITOCHONDRION ARE TRAFFICKED THROUGH THE GOLGI

In this study, we show that transport to the apicoplast of *Tg*TPx1/2, *Tg*SOD2 and *Tg*ACN/IRP, dually localised proteins of the apicoplast and the mitochondrion in *T. gondii* (Brydges, 2003; Pino et al., 2007), is diminished or perturbed in the presence of a C-terminal HDEL sequence. An HDEL sequence completely abolished the transport of *Tg*SOD2 to the apicoplast, while *Tg*TPx1/2 and *Tg*ACN/IRP were seen in the apicoplast with a background of a perinuclear spread as compared to the wild-type constructs. These observations are highly indicative of transport *via* the Golgi. Further, mutants of *Tg*TPx1/2 and *Tg*SOD2 that localised exclusively to the apicoplast were tested with an HDEL sequence. In this case, the HDEL sequence caused increased ER retention of these constructs along with apicoplast localisation, thus indicating the involvement of the Golgi in the transport of dually localised proteins to the apicoplast.

It is interesting to note that in the case of *Tg*TPx1/2, *Tg*ACN/IRP and the charged residue mutants *Tg*TPx1/2(R24A) and *Tg*SOD2(R12A), an HDEL sequence is unable to completely abolish apicoplast localisation and display retention in the ER alone. There could be two explanations for this phenotype. One is the fact that this is an overexpression system and overexpression of protein might overload the ER and cause “leakage”, as described in plants (Petruccelli et al., 2006; Abranches et al., 2008; De Meyer & Depicker, 2014). A second explanation could be that the transport of these proteins is not exclusively through the ER-Golgi route. These proteins, owing to their transit peptides, might be recognised by apicoplast-specific receptors in the ER and thus be trafficked to the apicoplast through both Golgi-dependent and -independent pathways.

Proteins that are trafficked through the ER and Golgi, travel *via* the machinery established for the classical secretory pathway. Protein cargo is delivered from the ER to the Golgi in COPII vesicles (Orci, Glick & Rothman, 1986; Barlowe et al., 1994). Proteins are loaded onto these vesicles upon recognition of ER exit motifs that are di-acidic (Nishimura & Balch, 1997), di-basic (Giraudo & Maccioni, 2003) or short stretches of hydrophobic amino acids (Kappeler et al., 1997). *Tg*TPx1/2, *Tg*SOD2 and *Tg*ACN have several of these motifs distributed throughout their primary sequences (data not shown). This distribution of motifs is seen across the proteins that travel through the Golgi-independent route as well (data not shown), indicating that these motifs may not be the factors differentiating this choice of pathway. Instead, these observations lead one to believe that the signal sequences of these two groups of proteins are different. The N-terminal signal sequences of dually localised proteins in *T. gondii* consist of signals that drive the proteins to both the mitochondrion and the apicoplast (Brydges, 2003; Mastud & Patankar, 2019). The similarity between the apicoplast transit peptide and a mitochondrial targeting sequence is that they are enriched in positively charged amino acids (Toursel et al., 2000; Truscott, Brandner & Pfanner, 2003; Brydges, 2003; Tonkin, Roos & McFadden, 2006), yet the N-terminus of a dually localised protein is an overlap of both these sequences and the resulting N-terminus is an ambiguous sequence capable of targeting the protein to both compartments (Mastud & Patankar, 2019). Hence, apicoplast proteins that use the Golgi-dependent trafficking route may have transit peptides with lower affinities for the ER receptors picking up cargo for delivery to the apicoplast.

### EVOLUTIONARY ORIGINS OF DUALLY LOCALISED PROTEINS DETERMINING TRANSPORT THROUGH TWO DISTINCT PATHWAYS

Signal sequences of apicoplast proteins and dually localised proteins are equipped with signals for the uptake of the protein into the ER before they are delivered to the apicoplast through a Golgi-dependent or - independent pathway. The puzzle still remains as to why proteins destined to the same organelles, despite their differences, adopt two distinct pathways to reach the apicoplast. A Golgi-dependent pathway is proposed to be a primitive pathway that the nascent endosymbiont utilised in the early stages of its integration into the host cell. It is hypothesised that the translocons for protein uptake and other endosymbiont proteins were initially trafficked through the secretory pathway of the host cell, until a separate pathway was established for transport into the plastid (Li & Chiu, 2010). Dually localised proteins in *T. gondii* have perhaps retained this ancient pathway, owing to their ambiguous targeting sequences. This hypothesis suggests that the dually localised proteins of the apicoplast may have evolutionary origins from the host genome, rather than from the apicoplast proteins that have been proposed to be derived from red algal ancestors.

Alternatively, these dually localised proteins might have been mitochondrial proteins that have gained a function at the apicoplast during the course of evolution. Gene transfer from the endosymbiotic organelles can often lead to multiple copies or loss of genes performing similar functions; this is accompanied by changes in the targeting sequences of these genes, favouring transport to a certain compartment (Martin, 2010). Mitochondrial proteins gaining a function at the apicoplast would have thus evolved an N-terminus with just enough signals to take it to both the apicoplast and the mitochondrion. Such ambiguous signal sequences direct dually localised proteins through a distinct pathway, as opposed to a Golgi-independent pathway of other apicoplast localised proteins.

Thus, there are two possible models for the evolution of dually localised proteins to the apicoplast and mitochondrion: first, the proteins have evolutionary origins in the apicoplast; they are closely related to plants, algae and cyanobacteria (Köhler et al., 1997; Blanchard & Hicks, 1999; McFadden, 2000; Fast et al., 2001; Funes et al., 2002; Harper & Keeling, 2003). These proteins have gained a function at the mitochondrion and the mitochondrial copy of the gene in the host genome was eventually lost. Second, the proteins are originally derived from mitochondrial evolution (proteobacteria)(Gray & Doolittle, 1982; Spencer, Schnare & Gray, 1984; Yang et al., 1985) and have gained a function at the apicoplast, whilst the apicoplast origin copy of the gene was lost over time. We observe that *Tg*TPx1/2 and *Tg*SOD2 seem to have originated from the mitochondrion, given their similarity to bacteria over plants and algae (Fig.4). *Tg*ACN, on the other hand, indicates close evolutionary relations to plants and algae distinct from other eukaryotic mitochondrial ACNs or bacteria, indicating apicoplast origins.

These results indicate that independently of the mitochondrial or plastid origins of the dually localised proteins, these proteins have still acquired similar signals and adopted similar trafficking pathways to the apicoplast. However, one cannot help but notice that the two proteins that have a mitochondrial/host origin are both involved in redox functions, leading to the speculation that a requirement for an anti-oxidant response in the apicoplast may have resulted in the selection of parasites that co-opted existing mitochondrial proteins to the apicoplast by using the available Golgi-dependent secretory pathway. The retention of this trafficking route till date may be due to other features, like a functional requirement of post-translational modifications at the Golgi.

The evolution of N-terminal signal sequences appears to be driven by the plasticity of signal sequences, enabling proteins derived from both the host and the endosymbiont to sample various compartments and acquire new functions (Martin, 2010). This might also lead to the selection of unique trafficking pathways influenced by strong positive selection of N-terminal signal sequences that have evolved to direct them into the right compartment efficiently.

In summary, this study proves the existence of two distinct protein trafficking pathways to the apicoplast. Apicoplast proteins seem to be trafficked to the plastid independent of the Golgi while dually localised proteins of the apicoplast and mitochondrion are trafficked through the Golgi to reach the apicoplast. Ambiguous signal sequences have not only led to dual localisation of proteins, but have also, perchance, exploited the evolution of two different pathways to the apicoplast.

## ACKNOWLEDGEMENTS

We acknowledge the Government of India, Department of Biotechnology (DBT) for the grant to SP that was utilized for funding this project (DBT/PR13546/BRB/10/1423/2015). We are grateful to the Government of India, Ministry of Human Resource Development (MHRD) and Indian Institute of Technology Bombay (IIT Bombay) for the Ph.D. fellowships given to AP and PM. We thank Dr. Dhanasekaran Shanmugam for generously sharing ACP antibodies and Prof. Kiran Kondabagil for his insights on evolutionary analysis. We appreciate Aishwarya Narayan’s timely and critical feedback on the manuscript. All imaging data were obtained at the confocal laser scanning microscopy facility funded by the Industrial Research and Consultancy Centre (IRCC) at IIT Bombay.

## AUTHORS’ CONTRIBUTIONS

AP, PM and SP conceived and designed the experiments. AP and PM performed the experiments. AP prepared the figures. AP and SP analysed the data and prepared the manuscript. AP, SP and PM reviewed drafts of the manuscripts.

