## Supplementary figure S1 for "Dually localized proteins found in both the apicoplast and mitochondrion utilize the Golgi-dependent pathway for apicoplast targeting in *Toxoplasma gondii*"

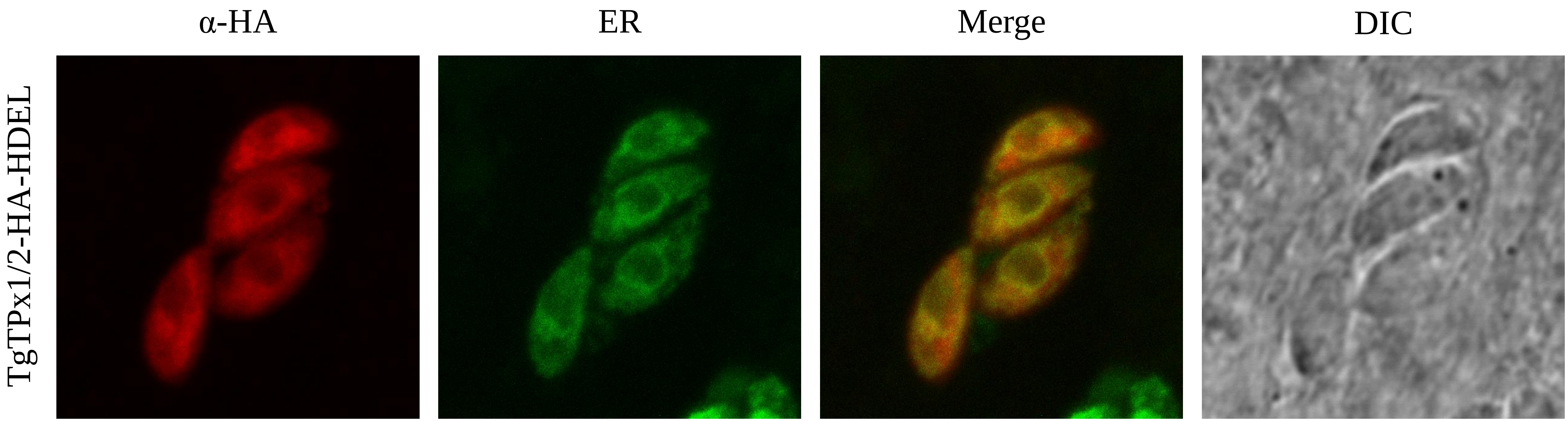


**S1**. Localisation of *Tg*TPx1/2-HA-HDEL with ER marker. *Tg*TPx1/2-HA-HDEL is stained with α-HA antibodies (in red) and the GFP tagged ER marker is shown in green. Colocalisation of the protein with the marker is indicated in yellow in the merged image.
